# Extreme Phenotypic Diversity in Operant Responding for an Intravenous Cocaine or Saline Infusion in the Hybrid Mouse Diversity Panel

**DOI:** 10.1101/2021.02.03.429584

**Authors:** Jared R. Bagley, Arshad H. Khan, Desmond J. Smith, James D. Jentsch

## Abstract

Cocaine self-administration is complexly determined trait, and a substantial proportion of individual differences in cocaine use is determined by genetic variation. Cocaine intravenous self-administration (IVSA) procedures in laboratory animals provide opportunities to prospectively investigate neurogenetic influences on the acquisition of voluntary cocaine use. Large and genetically diverse mouse populations, including the Hybrid Mouse Diversity Panel (HMDP), have been developed for forward genetic approaches that can reveal genetic variants that influence traits like cocaine IVSA. This population enables high resolution and well-powered genome wide association studies, as well as the discovery of genetic correlations. Here, we provide information on cocaine (or saline - as a control) IVSA in 65 strains of the HMDP. We found cocaine IVSA to be substantially heritable in this population, with strain-level intake ranging for near zero to >25 mg/kg/session. Though saline IVSA was also found to be heritable, a very modest genetic correlation between cocaine and saline IVSA indicates that operant responding for the cocaine reinforcer was influenced by a substantial proportion of unique genetic variants. These data indicate that the HMDP is suitable for forward genetic approaches for the analysis of cocaine IVSA, and this project has also led to the discovery of reference strains with extreme cocaine IVSA phenotypes, revealing them as polygenic models of risk and resilience to cocaine reinforcement. This is part of an ongoing effort to characterize genetic and genomic variation that moderates cocaine IVSA, which may, in turn, provide a more comprehensive understanding of cocaine risk genetics and neurobiology.

## Introduction

Cocaine use disorder is a psychiatric condition with a complex etiology, and genetic factors account for a substantial proportion of cocaine use disorder risk (Goldman et al., 2005; Kendler et al., 2000; Kendler & Prescott, 1998; van den Bree et al., 1998). Identification of specific risk genes, and the allelic variants that drive functional and expression variation to ultimately determine individual vulnerability will likely advance our understanding of the neurogenetic mechanisms that influence problematic cocaine use. Forward genetic approaches, in which unbiased, genome-wide scans can identify genetic variants that affect cocaine use disorder risk, have the potential to identify those risk alleles.

Forward genetic approaches can be employed both in study of human subjects and animal models for cocaine use and related traits. Human studies have had some success to date (Cabana-Domínguez et al., 2019; Gelernter et al., 2014; Marees et al., 2020); however, a large proportion of the genetic variants that contribute to heritable influence on cocaine use disorder remain undiscovered. Approaches in laboratory animal populations offer complementary methods with several key advantages. Advanced mouse populations allow for quantitative traits to be characterized prospectively, under controlled conditions. To date, forward genetic strategies have identified quantitative trait loci (QTL) and subsequent candidate genes for a number of cocaine-related behaviors. Some of these efforts have focused on cocaine-induced psychomotor activation; multiple QTL have been associated with cocaine-induced locomotion, with confirmation of some QTL by secondary mapping approaches (Boyle & Gill, 2001, 2009; Gill & Boyle, 2003; Jones et al., 1999; Kumar et al., 2013; Phillips, Huson, & McKinnon, 1998; Tolliver, Belknap, Woods, & Carney, 1994). Additional efforts have characterized measures of cocaine reward and reinforcement with success in mapping QTL and identifying candidate genes and genetic correlations among behavioral and physiological measures (Bagley, Szumlinski, et al., 2019; Bagley, Adams, Bozadjian, Bubalo, & Kippin, 2019; Cervantes et al., 2013; Dickson et al., 2014, 2015; Kippin et al., 2015; Roberts et al., 2018; Saul et al., 2020, 2020).

Cocaine intravenous self-administration (IVSA) is an assay of voluntary cocaine use and is thought to provide a model of cocaine use with predictive, face and construct validity (Belin-Rauscent et al., 2016; O’Connor et al., 2011). Advances in surgical techniques and devices have allowed for improved catheter patency rates that make characterization of large numbers of genetically diverse mice for cocaine IVSA feasible (Cervantes et al., 2013; Dickson et al., 2014, 2015; Roberts et al., 2018; Saul et al., 2020). Application of this assay to advanced mouse populations may enhance understanding of genetic factors unique to voluntary cocaine use. The BXD recombinant inbred strain panel was utilized to map cocaine IVSA QTL and genetic correlations (Dickson et al., 2015), and a similar study is underway using Collaborative Cross and Diversity Outbred mice (CC/DO) (Saul et al., 2020). Application of cocaine IVSA to additional mouse populations may offer additional opportunities to generate replicable and/or reproducible information on the genetics of cocaine use and related traits.

The Hybrid Mouse Diversity Panel (HMDP) is a large collection of both classic inbred strains and recombinant inbred strain panels derived from some of the those classical inbred strains (Bennett et al., 2010; Ghazalpour et al., 2012). This heterogenous panel provides a population with high genetic diversity and a large number of meiotic breakpoints that allows for relatively high-resolution QTL mapping. Inclusion of the recombinant inbred strains enhances power by adding additional replicates of alleles (Bennett et al., 2010; Ghazalpour et al., 2012). All mice of the HMDP are reference strains; the stability of the population allows for continued, cumulative study of traits, including characterization of transcript expression which may serve to greatly enhance success of genetic studies in current and future research. These key characteristics may be utilized to investigate the genetics of cocaine IVSA and related traits.

Here, we describe use of the HMDP to characterize cocaine IVSA for the first time. These efforts assessed the degree of heritability of behavioral traits exhibited during acquisition of cocaine IVSA and provide a database that will be utilized for future forward genetic approaches, including GWAS of behavioral outcomes and effects of cocaine IVSA on whole-transcriptome expression in key brain regions. A saline IVSA control group is included in every strain tested; this control may allow for enhanced precision in discriminating heritable cocaine reinforcement from other heritable factors that may influence IVSA behavior independently of cocaine.

Collectively, these data may serve to enhance understanding of the genetic factors that moderate cocaine self-administration.

## Materials and Methods

### Subjects

A total of 772 mice from sixty-five strains of the HMDP (mean age 11.2+/- 2.3 (SD) weeks) were selected for assessment of cocaine or saline IVSA (see Table 1). All animals received surgical implantation of chronic indwelling catheters by JAX Surgical Services (The Jackson Laboratory, Bar Harbor ME). Following at least one week of recovery, the animals were shipped to Binghamton University. When not being tested, all mice were maintained on *ad libitum* mouse chow (5L0D, Purina Lab Diet, MO USA) and water and were housed in polycarbonate cages (30 × 8 cm) with wood-chip bedding (SANI-CHIPS, NJ USA), a paper nestlet and a red polycarbonate hut, at a density of 1 mouse per cage. All procedures were approved by the Binghamton University Institutional Animal Care and Use Committee and conducted in accordance with the National Institute of Health Guide for Care and Use of Laboratory Animals (National Research Council (US) Committee for the Update of the Guide for the Care and Use of Laboratory Animals, 2011).

**Table 1.** Strain sample sizes for each strain and infusate group

| Classic Inbred Strain<br>(Strain ID) | Cocaine | Saline | BXD Strain<br>(Strain ID) | Cocaine | Saline |
| --- | --- | --- | --- | --- | --- |
| 129S1/SvImJ (2448) | 6 | 6 | BXD1/TyJ (36) | 6 | 7 |
| 129X1/SvJ (691) | 6 | 6 | BXD2/TyJ (75) | 6 | 5 |
| A/J (646) | 6 | 6 | BXD9/TyJ (105) | 6 | 5 |
| AKR/J (648) | 6 | 6 | BXD11/TyJ (12) | 5 | 5 |
| BALB/cByJ (1026) | 6 | 6 | BXD15/TyJ (95) | 5 | 6 |
| BALB/cJ (651) | 7 | 7 | BXD16/TyJ (13) | 5 | 6 |
| BPL/1J (3006) | 6 | 6 | BXD19/TyJ (10) | 5 | 6 |
| BTBRT+ltpr3tf/J (2282) | 7 | 6 | BXD21/TyJ (71) | 6 | 4 |
| C3H/HeJ (659) | 6 | 6 | BXD28/TyJ (47) | 5 | 6 |
| C3HeB/FeJ (658) | 6 | 6 | BXD31/TyJ (83) | 7 | 6 |
| C57BL/10J (665) | 6 | 7 | BXD32/TyJ (78) | 6 | 6 |
| C57BL/6J (664) | 7 | 6 | BXD34/TyJ (3223) | 5 | 6 |
| C57BLKS/J (662) | 6 | 6 | BXD38/TyJ (3227) | 6 | 6 |
| C57BR/cdJ (667) | 6 | 6 | BXD39/TyJ (3228) | 6 | 6 |
| C57L/J (668) | 6 | 8 | BXD40/TyJ (3229) | 5 | 6 |
| C58/J (669) | 6 | 6 | BXD42/TyJ (3230) | 6 | 6 |
| CBA/J (656) | 6 | 7 | BXD48a/RwwJ (7139) | 6 | 6 |
| DBA/1J (670) | 6 | 6 | BXD50/RwwJ (7099) | 6 | 6 |
| DBA/2J (671) | 6 | 6 | BXD55/RwwJ (7103) | 5 | 6 |
| FVB/NJ (1800) | 6 | 6 | BXD60/RwwJ (7105) | 4 | 6 |
| KK/Hij (2106) | 6 | 6 | BXD61/RwwJ (7106) | 6 | 6 |
| LP/J (676) | 6 | 6 | BXD62/RwwJ (7107) | 6 | 6 |
| MA/MyJ (677) | 6 | 6 | BXD63/RwwJ (7108) | 4 | 6 |
| MRL/MpJ (486) | 7 | 6 | BXD65/RwwJ (7110) | 6 | 6 |
| NOD/ShiLtJ (1976) | 6 | 7 | BXD68/RwwJ (7113) | 6 | 6 |
| NZB/BINJ (684) | 7 | 5 | BXD69/RwwJ (7114) | 6 | 4 |
| NZO/HILtJ (2105) | 5 | 7 | BXD70/RwwJ (7115) | 6 | 6 |
| NZW/LacJ (1058) | 6 | 6 | BXD73a/RwwJ (7124) | 6 | 6 |
| PL/J (680) | 6 | 6 | BXD75/RwwJ (7119) | 5 | 5 |
| SJL/J (686) | 7 | 6 | BXD77/RwwJ (7121) | 6 | 6 |
| SM/J (687) | 7 | 6 | BXD84/RwwJ (7127) | 6 | 6 |
|  |  |  | BXD86/RwwJ (7129) | 5 | 6 |
|  |  |  | BXD98/RwwJ (7141) | 6 | 6 |
|  |  |  | BXD100/RwwJ (7143) | 6 | 6 |

### Surgery

The JAX Surgical Services team anesthetized the animals with Tribromoethanol (Avertin, 400 mg/kg) injection and performed surgical procedures to implant chronic indwelling jugular catheters (catalog # CNC-2/3S-082109E/12, Access Technology, IL USA) and access button ports (1-VAB62SMBS/25, Instech, PA USA). Carprofen (5 mg/kg) was administered by subcutaneous injection pre- and post-surgically, and bupivacaine (0.1%: 2mg/kg) was applied topically before incision closure to manage any surgical pain.

### Catheter care and patency

Catheters were maintained with a flush of ~0.05 mL of sterile saline and by then filling the catheter with heparin lock solution (~0.01 mL volume, 500u/mL concentration, SAI Infusion Technologies, IL USA) at least once every 3 days during acclimation periods and daily during IVSA testing.

Catheter patency was confirmed by infusions of propofol (~0.02 mL volume, 10mg/mL concentration, Zoetis, NJ USA). Immediate but rapidly reversed loss of muscle tone indicated that the catheter was patent. This testing occurred once before the start of IVSA testing (3-4 days prior to testing) and again after the mouse completed the 10^th^ testing session. Any mice that failed the test were excluded from the experiment.

### Cocaine intravenous self-administration (IVSA)

All mice were assigned to one of two treatment groups (saline vs. cocaine IVSA), with half of the animals in each strain/sex randomly allocated to each (4-7 subjects in each strain/sex/group). All subjects were tested for acquisition of self-administration in 10 consecutive daily sessions (Fixed-ratio-1 schedule of reinforcement, 0 or 0.5 mg kg^-1^ body weight of cocaine per infusion) that ran until 65 infusions were earned or 2 h passed, whichever came first. Cocaine hydrochloride (Sigma Aldrich; St Louis MO) was dissolved in sterile saline at a concentration of 0.84 mg/mL to produce a freebase dose of 0.5 mg/kg/infusion. Infusion volume was 0.67 mL/kg/infusion for saline and cocaine groups. Testing occurred at the same time each day, during the light phase of a 12/12 h cycle. The animals were tested in Med Associates mouse self-administration chambers (55.69 x 38.1 x 35.56 cm, MED-307W-CT-D1, Med Associates, VT USA) that were fitted with 2 retractable ultrasensitive levers and that were housed within sound-attenuating cubicles. Assignment of the active infusion lever (right or left side of the box) was counterbalanced across strains/sex. Assignment of testing chamber was explicitly not random to minimize testing multiple mice from a given strain in the same chamber.

Test sessions began with the activation of the white noise and the illumination of 5 stimulus lights on the back of the chamber. No priming infusion(s) were delivered. When a subject actuated the active lever, an infusion was delivered and the house light flashed for 20 s. During this time-out period, contacts on the active lever were recorded but had no programmed consequence. Actuation of the inactive lever was also recorded and never had a programmed consequence.

### Data Analysis

Strain, infusate group, sex and IVSA session effects were assessed by ANOVA. Heritability was estimated by partial eta squared for the main effect of strain. Where appropriate, post-hoc comparisons were made, and Sidak p-value correction was utilized to correct for multiple testing. Infusions earned and lever discrimination (active versus inactive lever pressing) were the dependent variables of analysis. A p-value less than 0.05 was considered significant.

## Results

### Acquisition of IVSA

Mice from all 65 strains underwent 10 days of cocaine or saline IVSA, and the number of infusions earned per session (Figures 1A and 2) was assessed by mixed ANOVA with strain, sex, infusate group and session as factors. Infusions varied over sessions (main effect of session [F(5.3, 2675.9)=100.9, p<0.001), with ANOVA also revealing numerous higher-level interactions, including a session-strain-infusate group interaction [F(335.1,2675.9)=1.3, p<0.001], a session-strain-sex interaction [F(335.1,2675.9)=1.3, p<0.001], a session-infusate group interaction [F(1411.8,2675.9)=8.8, p<0.001] and a session-strain interaction [F(327.1, 2675.9)=2.0, p<0.001]).

**Figure 1.**
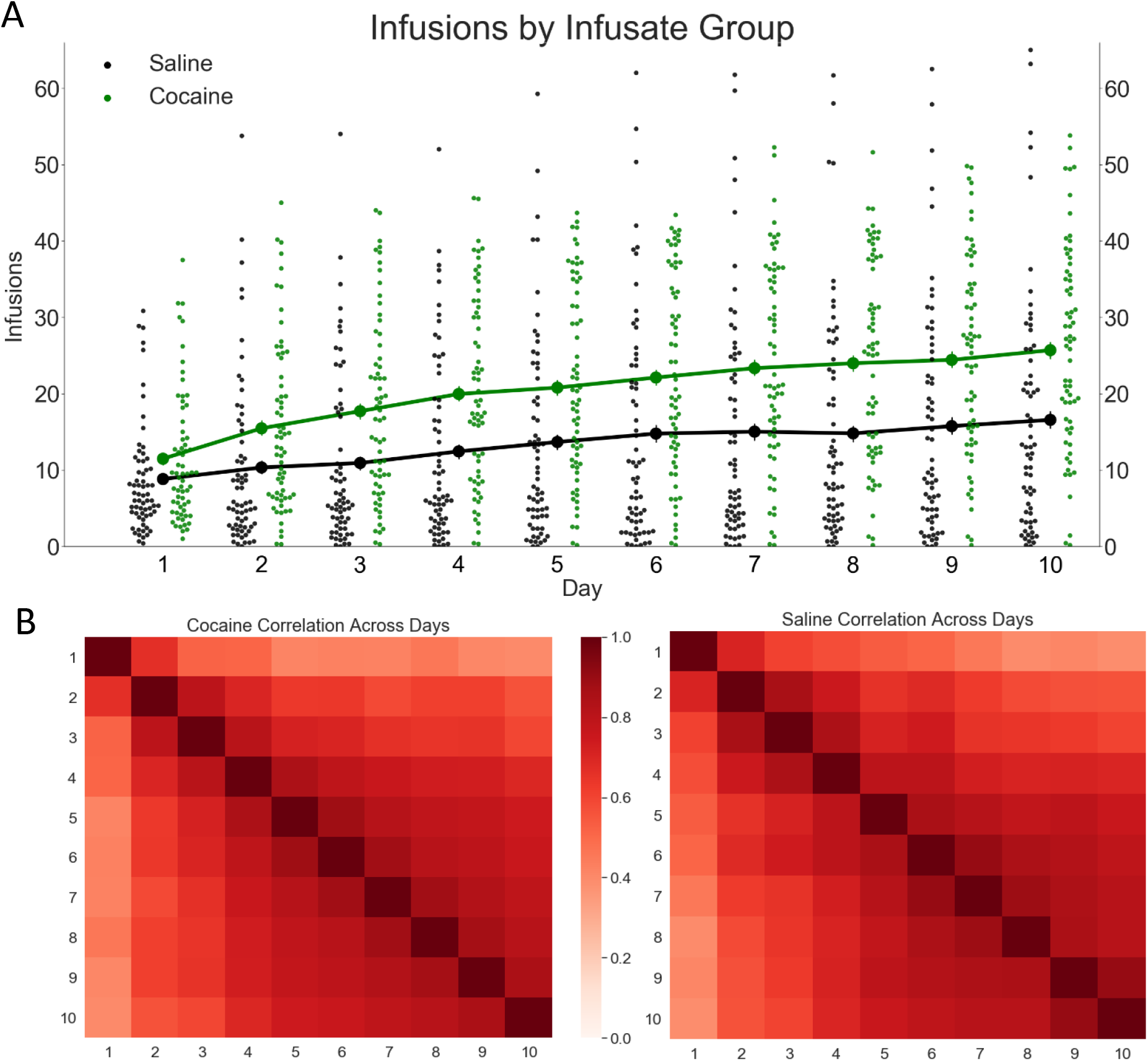
Cocaine infusions over 10 sessions, in 65 strains of the HMDP. A) Infusion means by infusate group, over 10 daily sessions (error bars =SEM, scatter points = strain means). Cocaine taking exceeded saline taking overall; however, IVSA of either infusate was highly strain-dependent. B) Correlation heatmap for correlation of infusions across sessions, for cocaine and saline groups. The magnitude of session-session correlation increased in later sessions, indicating stabilization of IVSA.

**Figure 2.**
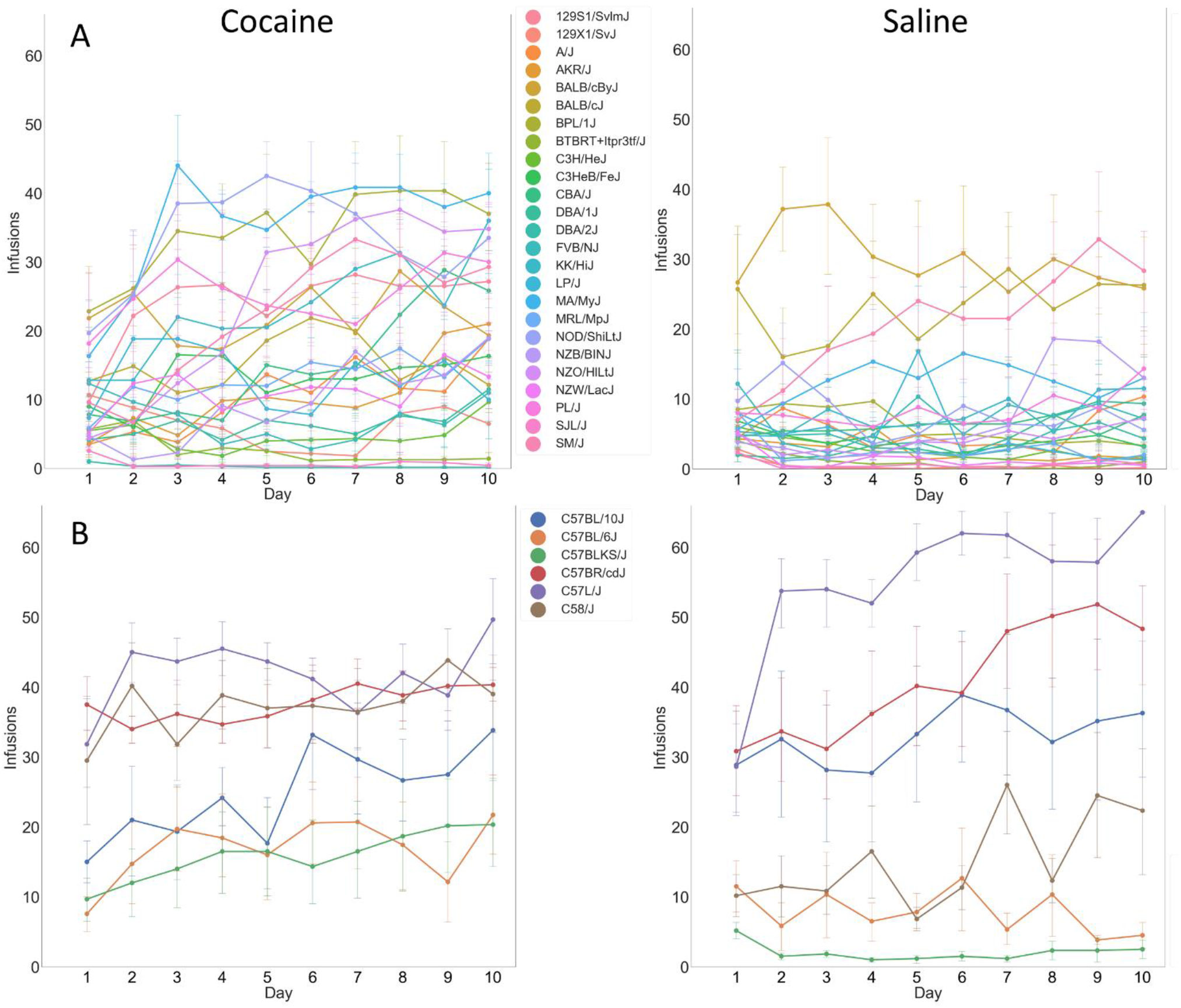
Infusions earned over 10 sessions in individual classic inbred strains (error bars=SEM). Strains are grouped into subcategories, based on relatedness, and graphed separately in order to improve clarity of the graphs. A) Classic inbred strains B) Strains of the C57 family.

Separate analyses within each infusate group revealed that both cocaine (main effect session [F(5.9,1480)=78,p<0.001]) and saline (main effect session [F(4.3,1123)=28.2, p<0.001]) groups increased the number of infusions earned as sessions progressed (Figures 1A and 2), and these effects were impacted by strain in both groups (session-strain interaction for the cocaine group [F(375.9,1480)=1.7, p<0.001] and for the saline group [F(277.7,1123)=1.7,p<0.001]). As expected, the cocaine group elicited more infusions overall (main effect of infusate group [F(1,511)=74.7,p<0.001]); however, differences between the two infusate groups were also impacted by strain (strain-infusate group interaction [F(64,511)=3.0, p<0.001]).

Analysis within infusate groups revealed significant broad sense heritability for cocaine IVSA (main effect strain [F(64,252)=4.7, p<0.001], H^2^=0.54) and saline IVSA (main effect strain [F(64,259)=7.1, p<0.001], H^2^=0.64). Sex interacted with strain to affect the number of infusions earned within the cocaine group [F(64,252)=1.9,p<0.001], but not the saline group, suggesting strain-dependent sex effects impact IVSA when cocaine is the reinforcer being earned.

### Post-Acquisition IVSA

Correlations between cocaine and saline infusions earned across all 10 test sessions revealed that day-to-day changes in infusions earned diminished as the number of sessions progressed (Figure 1B; test for difference between correlation r value from session 1 to 3 and session 8 to 10; z=10.5, p<0.001 and z=8.3,p<0.001 for cocaine and saline respectively). For this reason, infusions earned in sessions 8-10 were collapsed for each mouse and utilized as a measure of post-acquisition cocaine or saline IVSA.

Analyses of the sessions 8-10 phenotypes indicated that groups differed in infusions earned (main effect of infusate group [F(1,512)=74.5, p<0.001], with cocaine intake exceeding saline intake (Figures 3A). These differences were impacted by strain (strain-infusate group interaction [F(64,512)=2.9, p<0.001]). Analysis within each infusate group revealed significant broad-sense heritability for cocaine taking (main effect strain [F(64,253)=4.2, p<0.001], H^2^=0.52) and saline taking (main effect strain [F(64,259)=5.8, p<0.001], H^2^=0.59).

**Figure 3.**
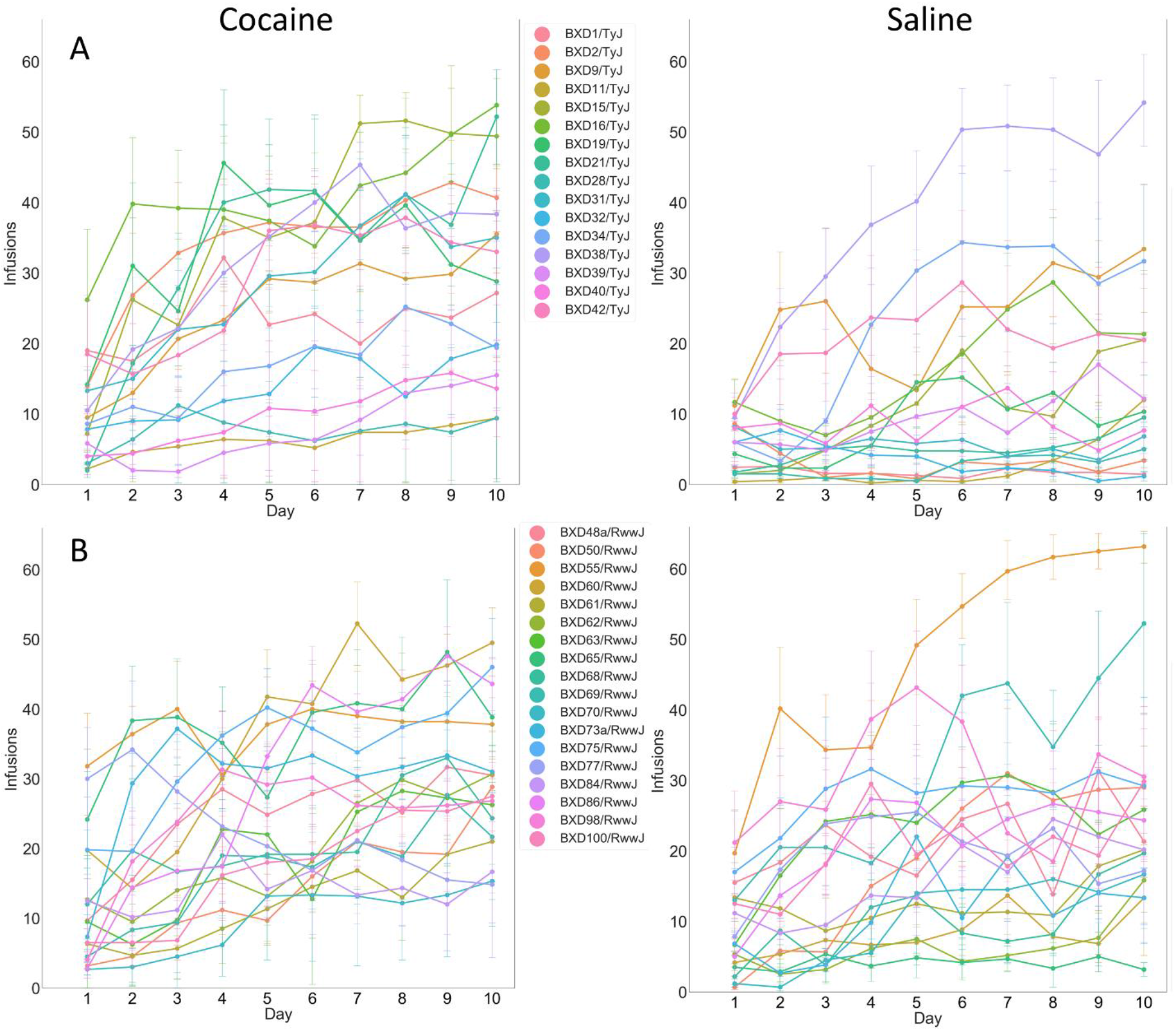
Infusions earned over 10 sessions in individual BXD strains (error bars=SEM). Strains are grouped into subcategories, based on relatedness, and graphed separately in order to improve clarity of the graphs. A) Taylor BXD strains B) Williams BXD strains.

Strains demonstrated a broad and continuous range of cocaine taking, with near zero intake at the low end and ~50 infusions (~25 mg/kg) per session at the high end (Figure 4A). The level of cocaine or saline self-administration in sessions 8-10 were assessed by one sample t-tests that compared each strain’s level of cocaine and saline taking to 0. Five of the 65 strains in the cocaine group did not, on average, earn a number of infusions that differed from 0 (Figure 4A). Similarly, 15 strains in the saline group did not demonstrate non-zero saline intake (Figure 5B).

**Figure 4.**
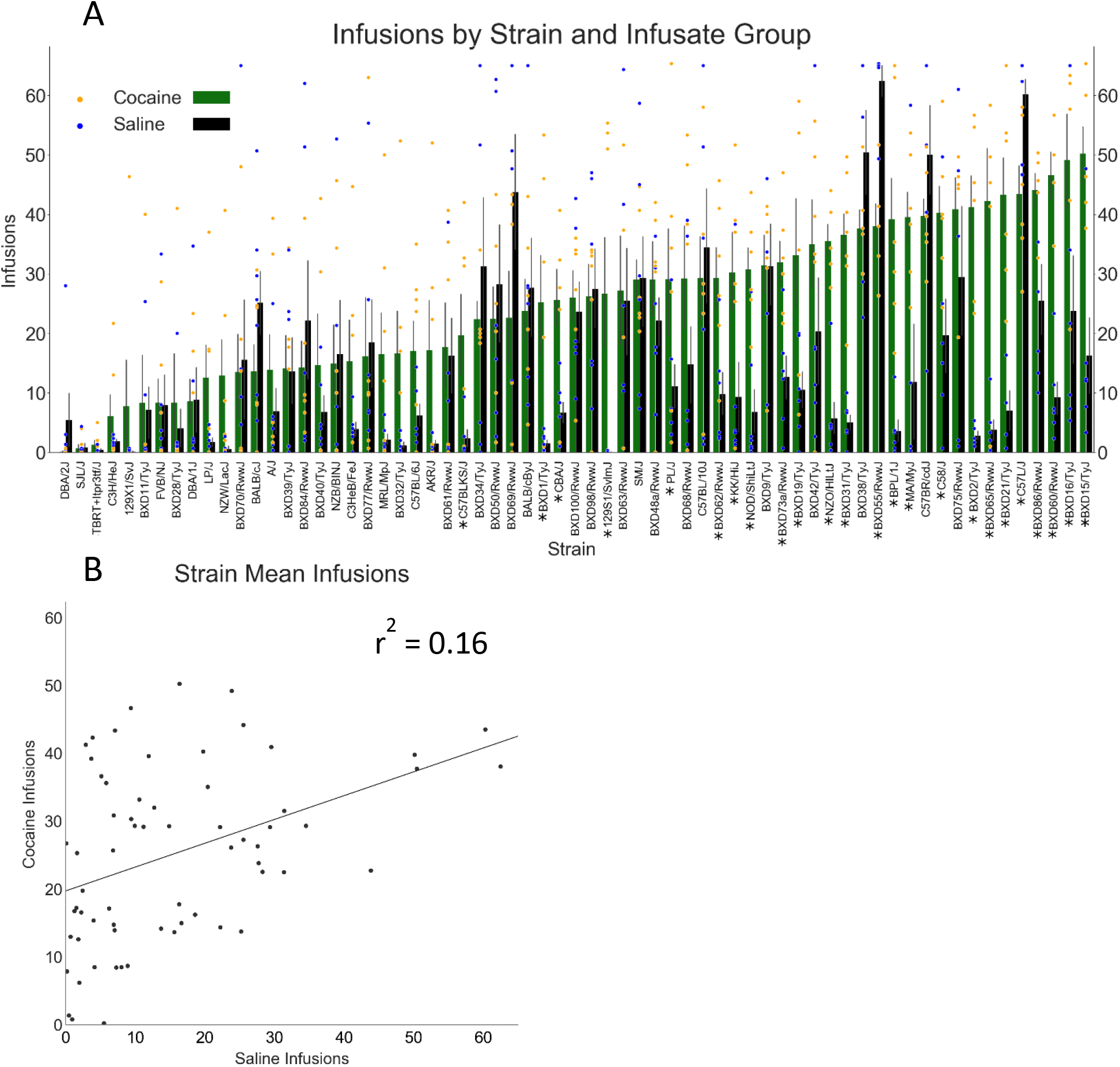
Cocaine and saline IVSA, averaged in the last three sessions. A) Strain means for cocaine and saline IVSA in the last 3 sessions. Both cocaine and saline demonstrated heritable variation, with a broad range of intake across strains (Error bars = SEM, scatter points = individual mouse values). A * below the strain name on the X-axis indicates that intake of cocaine or saline was significantly different for that strain. B) Scatterplot of strain means for cocaine and saline IVSA in the last three sessions. There was a modest genetic correlation between saline and cocaine taking.

**Figure 5.**
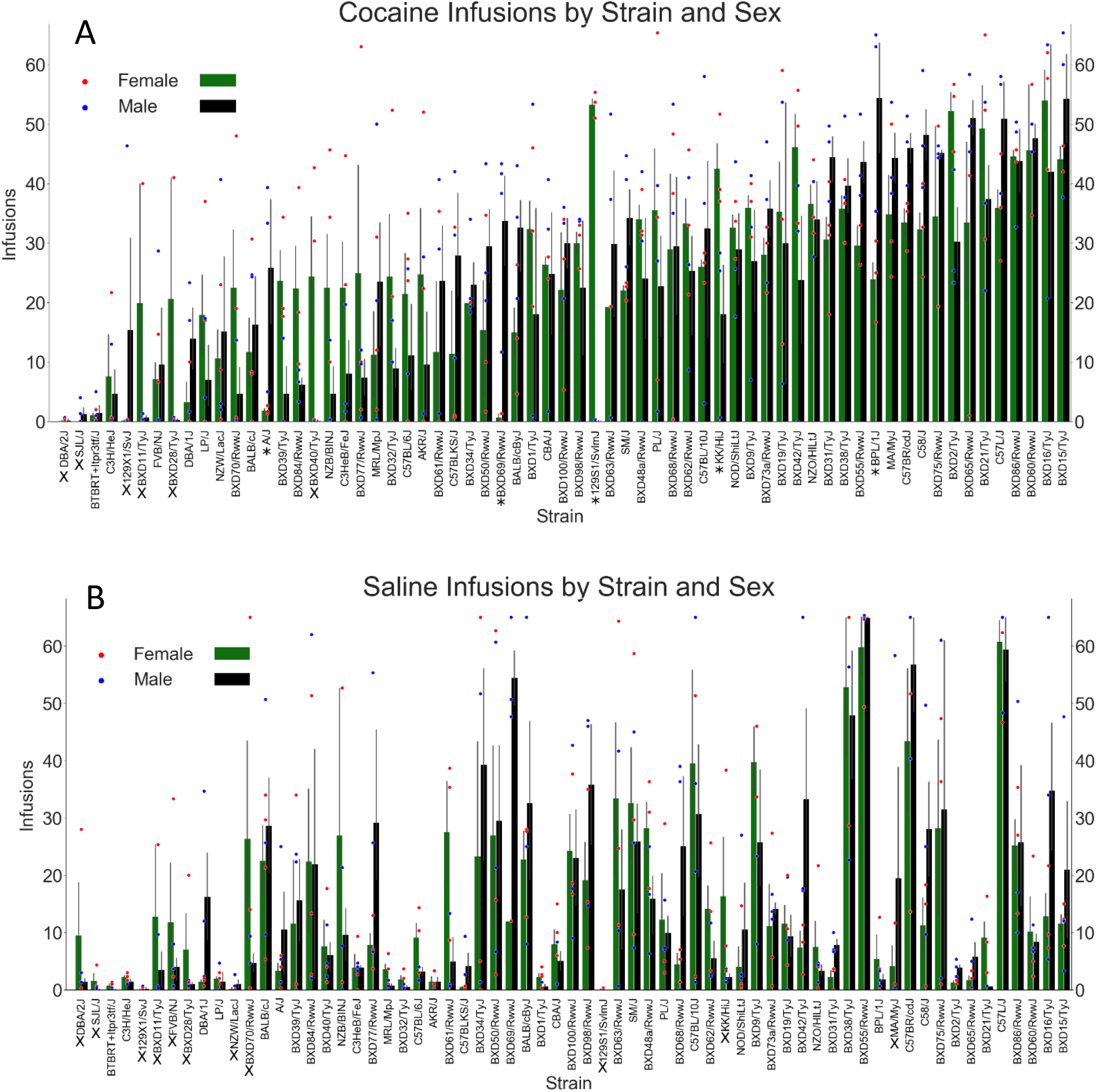
Cocaine and saline taking by strain and sex. A) Cocaine taking by strain and sex (error bars = SEM, scatter points = individual mouse values). Sex interacted with strain, indicating sex effects are moderated by genetic background. A * below the strain name indicates a significant sex difference within that strain. B) Saline taking by strain, ranked by levels of cocaine taking, and sex (error bars = SEM, scatter points = individual mouse values). No sex effects were discovered within the saline group. An X below the strain name, in graphs A and B, indicates that there was no significant difference between the number of infusions and zero for that strain.

Although strains demonstrated a broad range of saline taking, post-hoc comparisons of the two infusate groups, within strain, revealed that 24 strains demonstrated significantly different levels of cocaine and saline intake, with a majority of these strains (22 out of 24) demonstrating cocaine intake that exceeds saline intake (Figure 4A).

A strain-dependent sex effect was found for the session 8-10 average number of infusions earned (strain-sex interaction [F(64,512)=1.4, p=0.029]). Analysis within each infusate group revealed a strain-sex interaction in the cocaine group (Figure 5A) [F(64,253)=1.6,p=0.007] but not the saline group (Figure 5B). Post-hoc analyses for sex differences, within strain, in the cocaine group revealed that 5 strains demonstrated a significant sex difference. Females exhibited greater cocaine intake in 3 out 5 of these strains.

### Genetic correlation between cocaine and saline IVSA infusions

In order to assess the genetic correlation between cocaine and saline IVSA, a strain-level Pearson’s correlation was performed, examining the average number of infusions earned in sessions 8-10 on cocaine and saline groups. There was a significant, positive association between IVSA of cocaine and saline (r=0.40, p=0.001), revealing that a modest ~16% of the variance was shared between the two infusate group conditions (Figure 4B).

### Lever Preference

Preference for the active lever (active presses/total presses) was next assessed. The active lever triggers infusion of cocaine or saline and the active lever preference variable reveals whether lever pressing is random or reinforcer-directed. Because this variable often cannot be calculated early in training (when there are sessions with no presses on either lever) we focused on the later testing sessions (8-10).

The preference ratio was significantly greater in the cocaine group (main effect of infusate group [F(64,466)=80.2, p<0.001]), and lever preference differed by strain (main effect of strain [F(64,466)1.4, p=0.020]) (Figure 6A).

**Figure 6.**
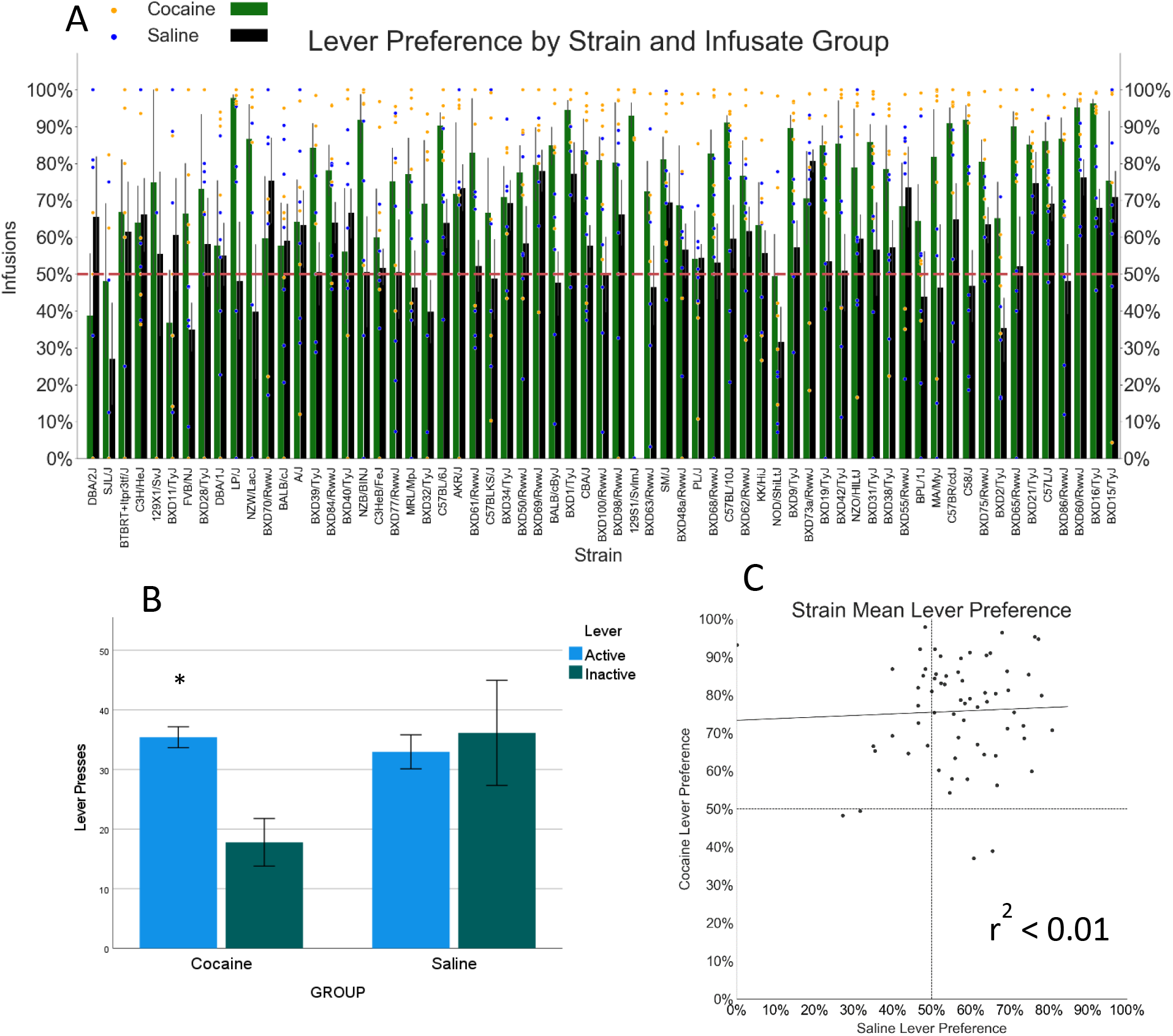
Lever discrimination in the last 3 sessions of IVSA. A) Active lever preference by strain and infusate group (error bars =SEM, scatterpoints = individual mouse values). Lever preference was greater in the cocaine group. B) Pressing on the active and inactive lever within the cocaine and saline groups. The cocaine group demonstrated greater active lever presses; the saline group did not. *=p<0.05 for difference between active and inactive lever pressing. C) Strain mean lever preference. No correlation between saline and cocaine lever preference was discovered.

A strain-level correlation was performed to assess genetic relationships between lever preference between cocaine and saline IVSA groups. No significant correlation was found (r=0.042, p=0.741) (Figure 6C).

Differences in lever pressing between the active and inactive lever, in sessions 8-10, were assessed by mixed ANOVA with lever as a within subject variable and strain, sex, and infusate group as between factors. A lever-infusate group interaction [F(1,512)=4.1,p=0.044] suggested lever discrimination varied in the two infusate groups (Figure 6B). Within group analysis revealed lever pressing differed by lever type in the cocaine group [F(1,253)=19.7, p<0.001], and this effect was impacted by strain and sex (lever-strain-sex interaction [F(64,253)=1.9, p<0.001] and a lever-strain interaction [F(64,253)1.7, p=0.002]). A strain-sex interaction [F(64,253)=2.0, p<0.001], main effect of strain [F(64,253)=2.5,p<0.001] and main effect of sex [F(1,253)=4.5,p<0.035] were also found within the cocaine group. No significant main effect of lever, or higher-level interactions involving lever, were found in the saline infusate group.

### Catheter patency and attrition

The mice were evaluated for catheter patency before and after the 10 d self-administration session, using infusion of propofol. The patency rate after the 10^th^ day of IVSA was 95.9%, reflecting a very high success rate for a protocol that demands high-quality small animal surgery, as well as animal care and husbandry. The overall attrition rate (patency failure, exclusion due to health issues or spontaneous death) was 8.4%. See **Table 1** for the numbers of mice by strain and group that have successfully completed IVSA.

## Discussion

The current research, which includes the largest published sample size for any IVSA study of rats or mice, reveals that cocaine IVSA is substantially heritable in this mouse genetic reference population, the HMDP, with more than half of all phenotypic variation being explained by strain. Strain-level infusions demonstrated a broad and continuous range of cocaine intake, from near zero to ~25 mg/kg per session. Given that the HMDP is suited for forward genetics analysis (Bennett et al., 2010; Ghazalpour et al., 2012), these data suggest that the results of an adequately powered final study (>100 strains, with n>=6 per strain per testing condition) may reveal genetic variants that associate with cocaine IVSA and lead to discovery of candidate genes that moderate cocaine use. Furthermore, individual strains can be identified as extreme responders and may serve as models of polygenic risk for high-level cocaine intake. These models may also be utilized to investigate the neurobiological mediators that underlie extreme differences in cocaine IVSA responding.

Mice received 10 daily sessions of cocaine or saline IVSA. We found that IVSA behavior changed over the 10 sessions in a strain-dependent manner. Many strains in the cocaine group increased their levels of intake over the sessions, consistent with experience-dependent acquisition of operant responding. Some strains did not increase intake to the same degree - or at all, and consequently, the variability in intake between strains increased as the training sessions progressed. Furthermore, correlations between sessions, at the individual mouse-level, were assessed, revealing that the magnitude of inter-session correlations increased as training progressed. In the final sessions (8-10), 60 of the strains demonstrated cocaine intake that was statistically significantly greater than zero. These results indicate acquisition and then stabilization of IVSA behavior, with both aspects being substantially moderated by genetic variation. Though a similar pattern of effects were observed in the saline infusate group, including both strain-dependent increases in responding across the sessions and greater inter-session stability in the final sessions, it is important to note that fewer strains (54) demonstrated a level of intake that was significantly different than zero. Notably, 22 of the strains exhibited significantly greater intake of cocaine compared to saline; given the modest number of mice per strain per infusate condition studied here, this is likely an underestimate of the real differences in cocaine and saline taking behavior.

Saline and cocaine IVSA are both heritable phenotypes in the HMDP; however, the genetic correlation between cocaine to saline IVSA was quite modest (~16% shared variance), with many strains demonstrating disparate cocaine and saline intake. Where intake differed within strain, cocaine IVSA was most often found to be greater than saline IVSA. These results suggest that some common genetic factors may influence self-administration of either infusate; however, those are less impactful than the unique genetic factors that influence intake of one infusate, but not the other. These saline data may be utilized to increase precision when measuring genetic influences on cocaine-specific reinforcement. Furthermore, these data suggest that some strains demonstrate equivalent cocaine and saline intake or greater intake of saline; with high levels of intake for both infusate types in some strains (e.g., C57L/J). Previous research found similar effects, with high, strain-dependent lever pressing in the absence of cocaine delivery and poor lever discrimination in the same strain during cocaine IVSA (Roberts et al., 2018). The degree to which high levels of cocaine taking in these strains reflects cocaine-seeking may be unclear and these outcomes highlight that inclusion of saline controls, particularly when investigating novel strains, may be prudent.

Lever pressing and infusions earned may the result of either associative or non-associative mechanisms in either infusate group. We found evidence that infusion-reinforced lever pressing occurred to a greater degree in the cocaine group. Within the cocaine group, we found significant preference for the active lever. An active lever preference was not found in saline group. In both infusate groups, activation of the active lever triggers a potentially perceptible infusion and a flashing house light; it is possible that the light stimulus serves as a reinforcer. Similar discrete light stimuli were sufficient to reinforce instrumental actions in the BXD population, and this trait demonstrated heritability (Dickson et al., 2015; Dickson & Mittleman, 2020). However, if the light stimulus served as an effective reinforcer in our procedure, a preference for the active lever that produces it would be expected but was not observed here. Alternatively, lever pressing in the saline infusate group may be the consequence of other phenotypes (e.g. increased exploratory motor activity in the test chamber, increased wakefulness, etc.). Taken together, these results suggest that there are some genetic variants that affect the potential to engage in lever pressing *per se*, with an additional set of variants affecting responding when cocaine is available.

Sex was found to moderate strain effects on responding in the cocaine, but not saline, group. Sex effects have been previously reported for mouse and rat cocaine IVSA (Algallal et al., 2020; Carroll et al., 2002; Davis et al., 2008; Jackson et al., 2005; Kerstetter et al., 2008; Lynch, 2008; Lynch et al., 2000; Lynch & Carroll, 2000; Roth & Carroll, 2004). Most often, females are found to have greater cocaine intake or faster acquisition. Here, we found that sex effects were highly strain-dependent and, in strains that demonstrate a sex effect, the direction of the effect is also strain-dependent. These results suggest that sex effects interact with genetic background to determine the presence and direction of the effect. This population may be utilized to identify the genetic variants that interact with sex to affect cocaine IVSA. Investigation of gene-sex hormone interactions and sex chromosome complement may be especially informative as both can have a large impact on cocaine taking and account for some of the sex differences in cocaine IVSA observed in rodent populations (Bagley, Adams, Bozadjian, Bubalo, Ploense, et al., 2019; Cummings et al., 2014; Hu & Becker, 2008; Kerstetter et al., 2012; Kippin et al., 2005; Larson et al., 2007; Lynch, 2008; Lynch et al., 2001; Martini et al., 2020; Perry et al., 2013; Ramoa et al., 2013; Segarra et al., 2010; Zhao & Becker, 2010). This approach may serve to further biological understanding of sex effects in cocaine use disorder.

These data demonstrate that cocaine IVSA is a highly heritable trait in the HMDP and that it is mostly genetically distinct from saline IVSA. In ongoing efforts, we will utilize the HMDP for forward genetic approaches and characterization whole-transcriptome, cocaine IVSA-induced changes in transcript expression. These complementary approaches will be utilized to identify candidate genes and gene networks that moderate cocaine IVSA. This research may contribute to a more comprehensive understanding of cocaine use disorder genetics and lead to more effective treatment techniques

## Acknowledgements

These studies were supported, in part, by Public Health Service grants U01-DA041602 (JRB, AMK, DJS, JDJ), T32-AA025606 (JDJ and JRB) and P50-DA039841(JDJ). We would like to thank Barbara Force for her assistance and technical support for this research. The authors have no conflicts of interest to declare.

## References

Algallal, H., Allain, F., Ndiaye, N. A., & Samaha, A.-N. (2020). Sex differences in cocaine self-administration behaviour under long access versus intermittent access conditions. Addiction Biology, 25(5), e12809. https://doi.org/10.1111/adb.12809

Bagley, J. R., Adams, J., Bozadjian, R. V., Bubalo, L., & Kippin, T. E. (2019). Strain differences in maternal neuroendocrine and behavioral responses to stress and the relation to offspring cocaine responsiveness. International Journal of Developmental Neuroscience, 78, 130–138. https://doi.org/10.1016/j.ijdevneu.2019.06.009

Bagley, J. R., Adams, J., Bozadjian, R. V., Bubalo, L., Ploense, K. L., & Kippin, T. E. (2019). Estradiol increases choice of cocaine over food in male rats. Physiology & Behavior, 203, 18–24. https://doi.org/10.1016/j.physbeh.2017.10.018

Bagley, J. R., Szumlinski, K. K., & Kippin, T. E. (2019). Discovery of early life stress interacting and sex-specific quantitative trait loci impacting cocaine responsiveness. British Journal of Pharmacology.

Belin-Rauscent, A., Fouyssac, M., Bonci, A., & Belin, D. (2016). How Preclinical Models Evolved to Resemble the Diagnostic Criteria of Drug Addiction. Biological Psychiatry, 79(1), 39–46. https://doi.org/10.1016/j.biopsych.2015.01.004

Bennett, B. J., Farber, C. R., Orozco, L., Kang, H. M., Ghazalpour, A., Siemers, N., Neubauer, M., Neuhaus, I., Yordanova, R., Guan, B., Truong, A., Yang, W., He, A., Kayne, P., Gargalovic, P., Kirchgessner, T., Pan, C., Castellani, L. W., Kostem, E.,… Lusis, A. J. (2010). A high-resolution association mapping panel for the dissection of complex traits in mice. Genome Research, 20(2), 281–290. https://doi.org/10.1101/gr.099234.109

Cabana-Domínguez, J., Shivalikanjli, A., Fernàndez-Castillo, N., & Cormand, B. (2019). Genome-wide association meta-analysis of cocaine dependence: Shared genetics with comorbid conditions. Progress in Neuro Psychopharmacology and Biological Psychiatry, 94, 109667. https://doi.org/10.1016/j.pnpbp.2019.109667

Carroll, M. E., Morgan, A. D., Lynch, W. J., Campbell, U. C., & Dess, N. K. (2002). Intravenous cocaine and heroin self-administration in rats selectively bred for differential saccharin intake: Phenotype and sex differences. Psychopharmacology, 161(3), 304–313. https://doi.org/10.1007/s00213-002-1030-5

Cervantes, M. C., Laughlin, R. E., & Jentsch, J. D. (2013). Cocaine self-administration behavior in inbred mouse lines segregating different capacities for inhibitory control. Psychopharmacology, 229(3), 515–525. https://doi.org/10.1007/s00213-013-3135-4

Cummings, J. A., Jagannathan, L., Jackson, L. R., & Becker, J. B. (2014). Sex differences in the effects of estradiol in the nucleus accumbens and striatum on the response to cocaine: Neurochemistry and behavior. Drug and Alcohol Dependence, 135, 22–28. https://doi.org/10.1016/j.drugalcdep.2013.09.009

Davis, B. A., Clinton, S. M., Akil, H., & Becker, J. B. (2008). The effects of novelty-seeking phenotypes and sex differences on acquisition of cocaine self-administration in selectively bred High-Responder and Low-Responder rats. Pharmacology Biochemistry and Behavior, 90(3), 331–338. https://doi.org/10.1016/j.pbb.2008.03.008

Dickson, P. E., Miller, M. M., Calton, M. A., Bubier, J. A., Cook, M. N., Goldowitz, D., Chesler, E. J., & Mittleman, G. (2015). Systems genetics of intravenous cocaine self-administration in the BXD recombinant inbred mouse panel. Psychopharmacology. https://doi.org/10.1007/s00213-015-4147-z

Dickson, P. E., & Mittleman, G. (2020). Stimulus Complexity and Mouse Strain Drive Escalation of Operant Sensation Seeking Within and Across Sessions in C57BL/6J and DBA/2J Mice. Frontiers in Behavioral Neuroscience, 13. https://doi.org/10.3389/fnbeh.2019.00286

Dickson, P. E., Ndukum, J., Wilcox, T., Clark, J., Roy, B., Zhang, L., Li, Y., Lin, D.-T., & Chesler, E. J. (2014). Association of novelty-related behaviors and intravenous cocaine self-administration in Diversity Outbred mice. Psychopharmacology, 232(6), 1011–1024. https://doi.org/10.1007/s00213-014-3737-5

Gelernter, J., Sherva, R., Koesterer, R., Almasy, L., Zhao, H., Kranzler, H. R., & Farrer, L. (2014). Genome-wide association study of cocaine dependence and related traits: FAM53B identified as a risk gene. Molecular Psychiatry, 19(6), 717–723. https://doi.org/10.1038/mp.2013.99

Ghazalpour, A., Rau, C. D., Farber, C. R., Bennett, B. J., Orozco, L. D., van Nas, A., Pan, C., Allayee, H., Beaven, S. W., Civelek, M., Davis, R. C., Drake, T. A., Friedman, R. A., Furlotte, N., Hui, S. T., Jentsch, J. D., Kostem, E., Kang, H. M., Kang, E. Y.,… LeBoeuf, R. C. (2012). Hybrid mouse diversity panel: A panel of inbred mouse strains suitable for analysis of complex genetic traits. Mammalian Genome, 23(9), 680–692. https://doi.org/10.1007/s00335-012-9411-5

Goldman, D., Oroszi, G., & Ducci, F. (2005). The genetics of addictions: Uncovering the genes. Nature Reviews Genetics, 6(7), 521–532. https://doi.org/10.1038/nrg1635

Hu, M., & Becker, J. B. (2008). Acquisition of cocaine self-administration in ovariectomized female rats: Effect of estradiol dose or chronic estradiol administration. Drug and Alcohol Dependence, 94(1-3), 56–62. https://doi.org/10.1016/j.drugalcdep.2007.10.005

Jackson, L. R., Robinson, T. E., & Becker, J. B. (2005). Sex Differences and Hormonal Influences on Acquisition of Cocaine Self-Administration in Rats. Neuropsychopharmacology, 31(1), 129–138. https://doi.org/10.1038/sj.npp.1300778

Kendler, K. S., Karkowski, L. M., Neale, M. C., & Prescott, C. A. (2000). Illicit psychoactive substance use, heavy use, abuse, and dependence in a US population-based sample of male twins. Archives of General Psychiatry, 57(3), 261–269. https://doi.org/10.1001/archpsyc.57.3.261

Kendler, K. S., & Prescott, C. A. (1998). Cocaine use, abuse and dependence in a population-based sample of female twins. The British Journal of Psychiatry: The Journal of Mental Science, 173, 345–350. https://doi.org/10.1192/bjp.173.4.345

Kerstetter, K. A., Aguilar, V. R., Parrish, A. B., & Kippin, T. E. (2008). Protracted time-dependent increases in cocaine-seeking behavior during cocaine withdrawal in female relative to male rats. Psychopharmacology, 198(1), 63–75. https://doi.org/10.1007/s00213-008-1089-8

Kerstetter, K. A., Ballis, M. A., Duffin-Lutgen, S., Carr, A. E., Behrens, A. M., & Kippin, T. E. (2012). Sex differences in selecting between food and cocaine reinforcement are mediated by estrogen. Neuropsychopharmacology: Official Publication of the American College of Neuropsychopharmacology, 37(12), 2605–2614. https://doi.org/10.1038/npp.2012.99

Kippin, T. E., Campbell, J. C., Ploense, K., Knight, C. P., & Bagley, J. (2015). Prenatal stress and adult drug-seeking behavior: Interactions with genes and relation to nondrug-related behavior. Advances in Neurobiology, 10, 75–100. https://doi.org/10.1007/978-1-4939-1372-5_5

Kippin, T. E., Fuchs, R. A., Mehta, R. H., Case, J. M., Parker, M. P., Bimonte-Nelson, H. A., & See, R. E. (2005). Potentiation of cocaine-primed reinstatement of drug seeking in female rats during estrus. Psychopharmacology, 182(2), 245–252. https://doi.org/10.1007/s00213-005-0071-y

Larson, E. B., Anker, J. J., Gliddon, L. A., Fons, K. S., & Carroll, M. E. (2007). Effects of estrogen and progesterone on the escalation of cocaine self-administration in female rats during extended access. Experimental and Clinical Psychopharmacology, 15(5), 461–471. https://doi.org/10.1037/1064-1297.15.5.461

Lynch, W. J. (2008). Acquisition and maintenance of cocaine self-administration in adolescent rats: Effects of sex and gonadal hormones. Psychopharmacology, 197(2), 237–246. https://doi.org/10.1007/s00213-007-1028-0

Lynch, W. J., Arizzi, M. N., & Carroll, M. E. (2000). Effects of sex and the estrous cycle on regulation of intravenously self-administered cocaine in rats. Psychopharmacology, 152(2), 132–139. https://doi.org/10.1007/s002130000488

Lynch, W. J., & Carroll, M. E. (2000). Reinstatement of cocaine self-administration in rats: Sex differences. Psychopharmacology, 148(2), 196–200. https://doi.org/10.1007/s002130050042

Lynch, W. J., Roth, M. E., Mickelberg, J. L., & Carroll, M. E. (2001). Role of estrogen in the acquisition of intravenously self-administered cocaine in female rats. Pharmacology Biochemistry and Behavior, 68(4), 641–646. https://doi.org/10.1016/S0091-3057(01)00455-5

Marees, A. T., Gamazon, E. R., Gerring, Z., Vorspan, F., Fingal, J., van den Brink, W., Smit, D. J. A., Verweij, K. J. H., Kranzler, H. R., Sherva, R., Farrer, L., Gelernter, J., & Derks, E. M. (2020). Post-GWAS analysis of six substance use traits improves the identification and functional interpretation of genetic risk loci. Drug and Alcohol Dependence, 206, 107703. https://doi.org/10.1016/j.drugalcdep.2019.107703

Martini, M., Irvin, J. W., Lee, C. G., Lynch, W. J., & Rissman, E. F. (2020). Sex chromosome complement influences vulnerability to cocaine in mice. Hormones and Behavior, 125, 104821. https://doi.org/10.1016/j.yhbeh.2020.104821

O’Connor, E. C., Chapman, K., Butler, P., & Mead, A. N. (2011). The predictive validity of the rat self-administration model for abuse liability. Neuroscience & Biobehavioral Reviews, 35(3), 912–938. https://doi.org/10.1016/j.neubiorev.2010.10.012

Perry, A. N., Westenbroek, C., & Becker, J. B. (2013). Impact of pubertal and adult estradiol treatments on cocaine self-administration. Hormones and Behavior, 64(4). https://doi.org/10.1016/j.yhbeh.2013.08.007

Ramoa, C. P., Doyle, S. E., Naim, D. W., & Lynch, W. J. (2013). Estradiol as a Mechanism for Sex Differences in the Development of an Addicted Phenotype following Extended Access Cocaine Self-Administration. Neuropsychopharmacology, 38(9), 1698–1705. https://doi.org/10.1038/npp.2013.68

Roberts, A. J., Casal, L., Huitron-Resendiz, S., Thompson, T., & Tarantino, L. M. (2018). Intravenous cocaine selfa-dministration in a panel of inbred mouse strains differing in acute locomotor sensitivity to cocaine. Psychopharmacology, 235(4), 1179–1189. https://doi.org/10.1007/s00213-018-4834-7

Roth, M. E., & Carroll, M. E. (2004). Sex differences in the escalation of intravenous cocaine intake following long-or short-access to cocaine self-administration. Pharmacology Biochemistry and Behavior, 78(2), 199–207. https://doi.org/10.1016/j.pbb.2004.03.018

Saul, M. C., Bagley, J. R., Bailey, L. S., Datta, U., Dickson, P. E., Dodd, R., Gagnon, L. H., Hugett, S. B., Kimble, V. M., Leonardo, M., Kim, S.-M., Olson, A., Roy, T., Schoenrock, S. A., Wilcox, T., Jentsch, J. D., Logan, R. W., McClung, C. A., Palmer, R. H. C.,… Chesler, E. J. (2020). Consideration of genetic and sex effects in mice enhances consilience with human addiction studies. BioRxiv, 2020.02.14.949784. https://doi.org/10.1101/2020.02.14.949784

Segarra, A. C., Agosto-Rivera, J. L., Febo, M., Lugo-Escobar, N., Menendez-Delmestre, R., Puig-Ramos, A., & Torres-Diaz, Y. M. (2010). Estradiol: A key biological substrate mediating the response to cocaine in female rats. Hormones and Behavior, 58(1), 33–43. https://doi.org/10.1016/j.yhbeh.2009.12.003

van den Bree, M. B., Johnson, E. O., Neale, M. C., & Pickens, R. W. (1998). Genetic and environmental influences on drug use and abuse/dependence in male and female twins. Drug and Alcohol Dependence, 52(3), 231–241. https://doi.org/10.1016/s0376-8716(98)00101-x

Zhao, W., & Becker, J. B. (2010). Sensitization enhances acquisition of cocaine self-administration in female rats: Estradiol further enhances cocaine intake after acquisition. Hormones and Behavior, 58(1), 8–12. https://doi.org/10.1016/j.yhbeh.2009.09.005

